# Cortical bursts provide spatio-temporal frames for the large-scale phase synchrony emerging across the birth transition

**DOI:** 10.64898/2026.10.28.762417

**Authors:** Sebastian König, Pauliina Yrjölä, Sampsa Vanhatalo, Anton Tokariev

## Abstract

Structural corticocortical connectivity grows rapidly around birth, but how it establishes the large-scale neuronal interactions needed for lifelong neurocognitive performance remains unclear. Using repeated high-density EEG from 233 preterm-born infants recorded at 33–45 weeks postmenstrual age, we characterized the development of three phase-based interaction modes with distinct temporal relationships to cortical bursts: burst-based, burst-framed, and general phase-based connectivity. Burst-framed connectivity was quantified with a novel metric, nested phase-phase correlation. All three modes declined globally with age but differed spatially: burst-based connectivity became more confined, burst-framed connectivity reorganized towards sensorimotor-association connectivity, whereas general phase-based connectivity retained a stable modular organization. General phase-based connectivity was initially concentrated along burst-framed connections, but this coupling weakened with age. Its greater persistence around term age was associated with poorer neurocognitive outcomes, suggesting delayed network maturation. These findings suggest that bursting may transiently scaffold the maturation of large-scale corticocortical phase synchrony across the birth transition.

## Introduction

Higher-order brain functions rely on precisely coordinated neuronal activity across large-scale networks (Bressler and Menon, 2010; Engel et al., 2013; Uddin et al., 2023; van Es et al., 2025; Vinck et al., 2023), where concurrent operations are carried out within shared structural networks by dynamic, spatially overlapping patterns of activity (Honey et al., 2009; Li et al., 2023; Spampinato et al., 2026). How these networks of neuronal activity emerge during development is central to understanding how higher-order brain functions arise.(França et al., 2024; Keunen et al., 2017; Tokariev et al., 2016b; Yrjölä et al., 2024). The prenatal structural development of the long-range core brain networks is essentially hard-wired (Chen et al., 2024; Dubois et al., 2015), relying on genetically determined region-specific molecular cues (Chédotal and Richards, 2010; Zhou et al., 2024) that guide development of excessive and unspecific structural connectivity (Innocenti and Price, 2005; Wolterhoff and Hiesinger, 2024).

Studies on sensory systems have shown how the coarse, primary wiring is subsequently organized in an activity-dependent manner by the temporally and spatially patterned endogenous activity within each sensory system, yielding their well-known somatotopic, retinotopic and tonotopic organizations, respectively (Cline et al., 2023; Katz and Shatz, 1996; Kirkby et al., 2013; Martini et al., 2021). It is intuitively clear, however, that the straightforward uni-sensory training mechanisms (Cline et al., 2023; Katz and Shatz, 1996; Kirkby et al., 2013) cannot train the brain-wide corticocortical ensembles. Numerous experimental, clinical, and simulation studies have also demonstrated remarkable functional resilience of higher-order brain functions to early anatomically circumscribed lesions (Aerts et al., 2016; Craig et al., 2020; Kirton et al., 2021; Newport et al., 2022). Taken these considerations together, there must be an endogenous and anatomically distributed mechanism to drive the early developmental training of brain-wide corticocortical activity networks.

Recent neuroscience literature suggests a candidate for such network training: the early neuronal ensembles exhibit an endogenous, spatio-temporally patterned activity, which emerges spontaneously during early phases of neuronal network development (Luhmann et al., 2022; Martini et al., 2021), and it can readily span across brain’s functional and hierarchic divisions (Khazaei et al., 2023; Luhmann et al., 2016; Tokariev et al., 2016b; Yrjölä et al., 2024). In the human infant, late-fetal cortical neural activity can be assessed non-invasively using scalp electroencephalography (EEG) in preterm-born infants, because their cortical activity follows conceptional age rather than postnatal age (Pavlidis et al., 2025; Stevenson et al., 2020). The hallmark of preterm EEG is intermittent, multifrequency bursting (a.k.a. spontaneous activity transients; (Luhmann et al., 2022; Molnár et al., 2020; Vanhatalo and Kaila, 2006; Wallois et al., 2021)), which exhibits initially global and unspecific correlations with developmental convergence to spatially and functionally confined networks by the end of pregnancy (Omidvarnia et al., 2014; Tokariev et al., 2016b; Wallois et al., 2021; Yrjölä et al., 2024), providing a seemingly clear candidate mechanism for training the early cortical networks

In the present study, we set out to characterize development of the endogenous cortical activity mechanisms in human infants that could account for the late fetal emergence of precisely coordinated neuronal activity in the brain-wide corticocortical networks (Tokariev et al., 2016b; Yrjölä et al., 2024). We hypothesized that the EEG bursts provide specific spatio-temporal windows for the early corticocortical correlations, with gradually changing connectivity mechanisms across the birth transition. To test this idea, we postulated an incrementally specific coordination in corticocortical activity: First, the burst-related activity in one cortical region is initially linked to co-occurring fast oscillatory activities across distant cortical regions (burst-based connectivity), which can be measured as distant phase-amplitude coupling (d-PAC, (Siebenhühner et al., 2020; Yrjölä et al., 2024)). Second, corticocortical phase synchrony of faster oscillatory activity is preferentially expressed during the bursts that provide temporal windows between cortical regions (burst-framed connectivity), which can be measured using a novel metric, nested phase-phase correlation (NPPC). Third, alongside the late-fetal decline of cortical bursting (Vanhatalo et al., 2005; Wallois et al., 2021), the cortical phase–phase correlation (PPC) networks become burst-independent and exhibit spatially consistent brain-wide constellations (Tokariev et al., 2019; Yrjölä et al., 2024). Finally, we postulate that these developmental passages align anatomically with the hierarchical sensorimotor-association axis (S-A) that underpin global brain functions throughout later life (Larsen et al., 2023; Sydnor et al., 2021), while they also link developmentally to the infants’ later emerging neurocognitive performance.

## Results

The study overview and the mechanistic hypotheses underlying the present EEG metrics are presented in Figure 1. Altogether, 498 EEG recordings from 233 preterm infants (1–4 per infant) were acquired during daytime sleep at 33–45 weeks of conceptional age (term birth is around 40 weeks) using a 124-channel high-density EEG system. Five-minute EEG epochs of the two cardinal neonatal sleep states, active and quiet sleep, were source-reconstructed into cortical activity across 58 parcels (Tokariev et al., 2019; Yrjölä et al., 2024) spanning sensorimotor (S) and association (A) regions, and filtered into four frequency bands: low delta (0.4–1.5 Hz), theta (4–8 Hz), alpha (8–13 Hz), and beta (13–22 Hz). In both sleep states, three complementary corticocortical connectivity modes were computed between all pairs of cortical parcel signals: burst-based (d-PAC), burst-framed (NPPC), and burst-independent connectivity (PPC), with low delta serving as the low-frequency component for d-PAC and NPPC (Tokariev et al., 2022; Yrjölä et al., 2024), while connectivity was assessed in theta, alpha, and beta. The developmental changes in connectivity modes were assessed at global and spatial network levels, as well as in relation to the hierarchical sensorimotor-association axis (S-A) (Ball et al., 2020; Larsen et al., 2023). In particular, we studied whether the most temporally precise communication mode, PPC, is emphasized within NPPC networks, and whether it is linked to later neurobehavior assessed using standardized Bayley Scales scores (Bayley, 2006).

**Figure 1.**
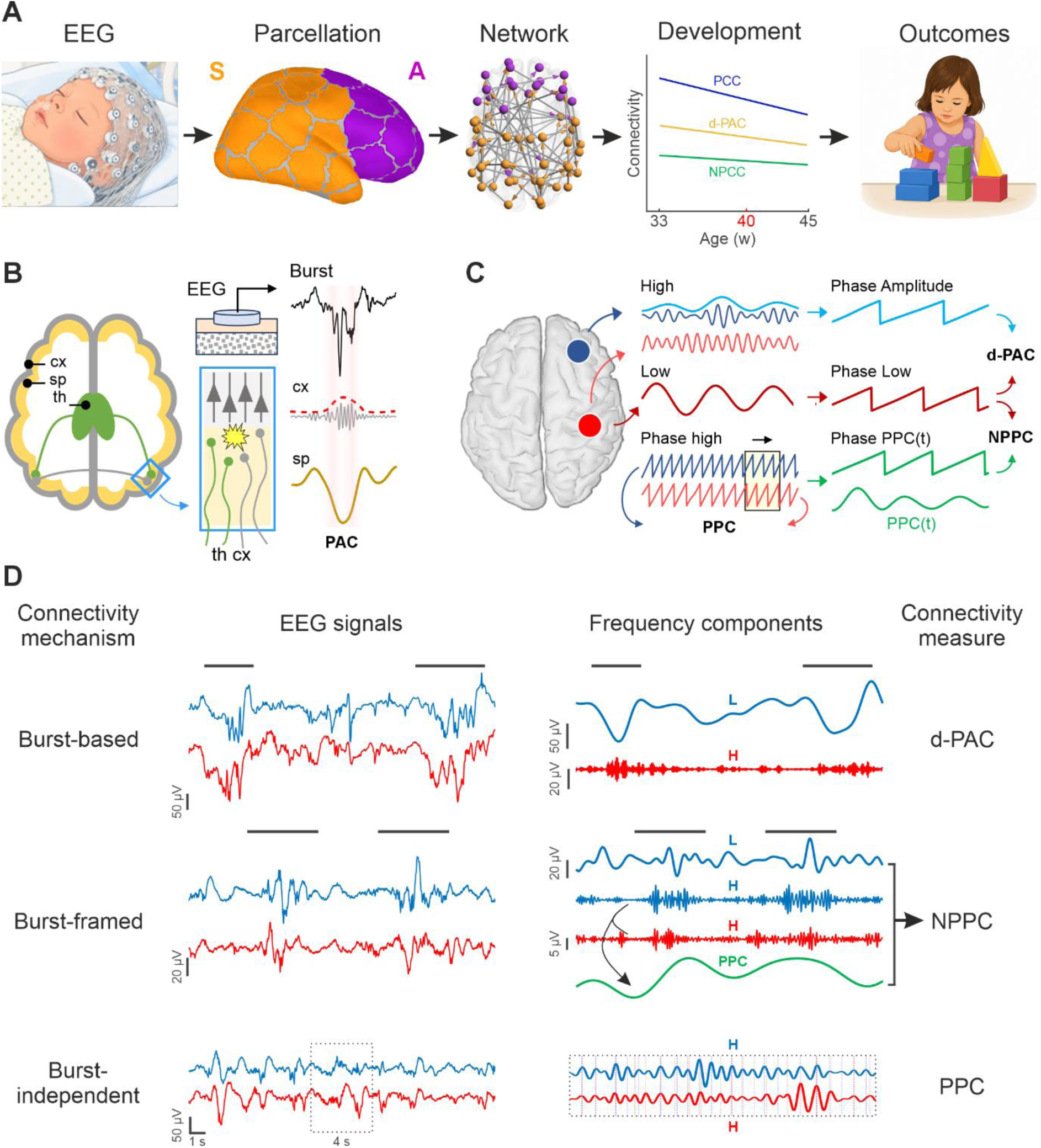
Study design and mechanistic hypothesis. **A**, Overview of the analytical flow. High-density EEG recordings from 233 preterm infants between 33 and 45 weeks of conceptional age were analysed (498 recordings overall). EEG signals were reconstructed as cortical activity across 58 parcels spanning the sensorimotor-association (S-A) axis. Interregional connectivity reflecting distinct interaction modes (see C) provided mode-specific functional networks. Developmental changes in global strength and organization were compared across these modes. Finally, network findings were associated with neurodevelopmental outcomes at 18 months. **B**, Presumed physiological underpinnings of cortical bursting, involving transient interactions between cortex (cx) and subplate (sp), with potential contributions from thalamocortical (th-cx) or corticocortical (cx-cx) inputs. The complex multifrequency EEG burst waveform is characterized by phase-amplitude coupling (PAC) between low-frequency, high-amplitude activity from deeper cortical and subplate structures and higher-frequency intracortical activity. Co-occurring bursts across distinct cortical regions may provide temporal windows that facilitate entrainment of higher-frequency phase-based corticocortical networks. **C**, Schematic computation of distant PAC (d-PAC), nested phase-phase correlation (NPPC), and phase-phase correlation (PPC). d-PAC is calculated between the phase of the low-frequency signal and the phase of the amplitude envelope of high-frequency activity originating from another cortical location. For NPPC, the time-resolved phase-phase connectivity, PPC(t), is computed from the high-frequency phases at two distinct cortical locations, and the phase of PPC(t) is coupled to the phase of the low-frequency signal from one of these locations. PPC quantifies phase locking between high-frequency signals from two cortical regions. **D**, Overview of the connectivity measures used in the present study, linking each connectivity mode to an example pair of infant EEG signals, its analytical manipulation, and the resulting connectivity measures: d-PAC, NPPC, and PPC. Frequency components (low, L; high, H) of the corresponding EEG signals are shown in the same colours.

### Burst-based connectivity declines and becomes increasingly organized within sensorimotor and association regions

Global levels of burst-based connectivity (d-PAC, top panel in Fig. 1C) showed a robust decrease with age (R^2^ = 0.41; P < 0.001; Fig. 2A). The spatial organization of burst-based networks changed clearly both between and within S and A regions (Fig. 2B). In the younger infants (33–41 weeks), d-PAC networks comprised predominantly the within-S connections with developmentally stable density (3.2–3.7%), whereas density of the A→S connections declined over time (from 3.6% to 1.4%). At post-term age (41–45 weeks), however, spatial organization shifted markedly towards within-A connections (9.5%), particularly involving frontal regions. These spatial patterns were robust across alternative thresholds (Supplementary Fig. 1A). A pronounced developmental shift was also observed in the distribution of connection lengths from long-range dominance in the preterm infants to short-range dominance after full-term age. Fig. 2C). These findings were largely consistent across sleep states and frequency bands (for detailed results, see Supplementary Fig. 2). Taken together, burst-based connectivity declines globally and shifts towards short-range connections within sensorimotor and association regions, with a particular increase in frontal connectivity after term age.

**Figure 2.**
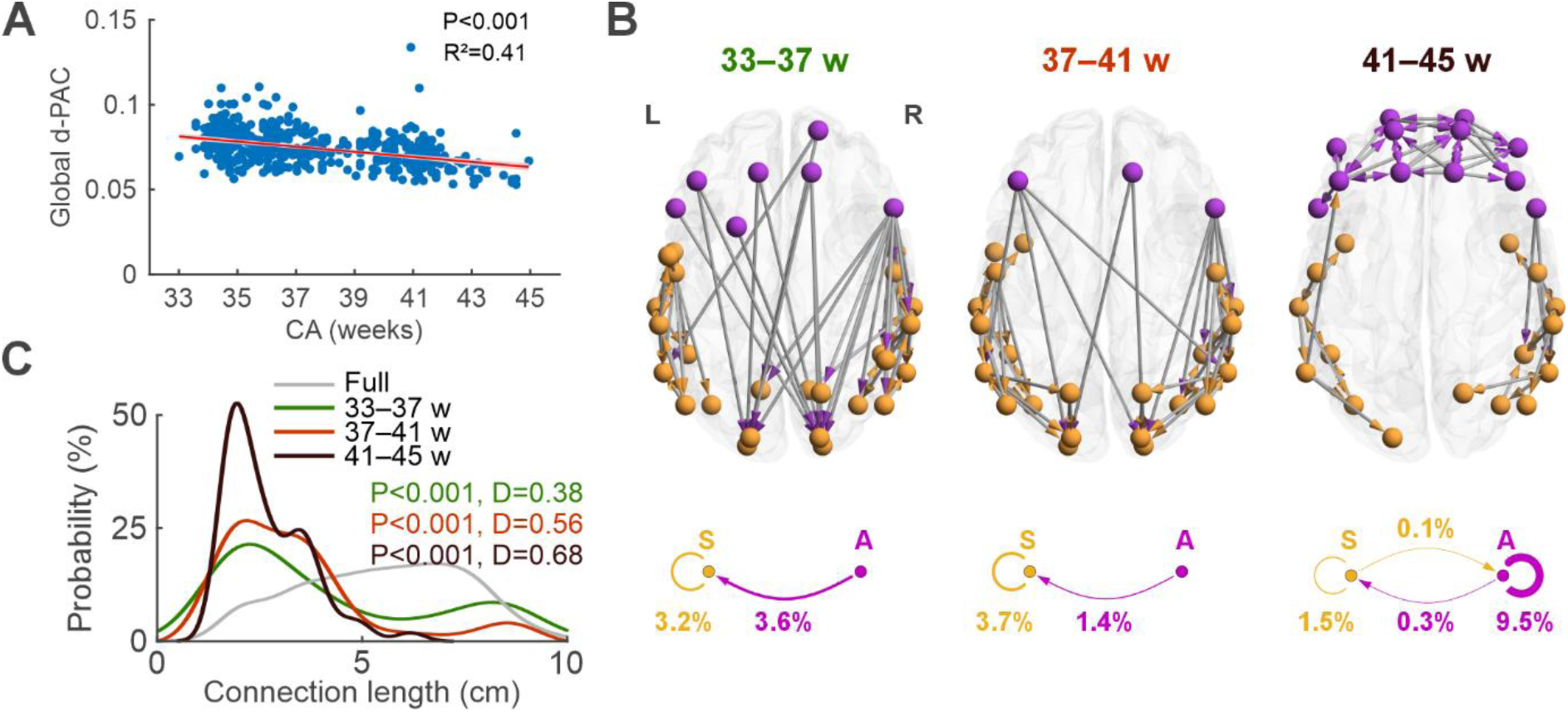
Development of burst-based connectivity (d-PAC). **A**, Developmental decline in global d-PAC strength. Each point represents one EEG recording, and the red line shows the fitted LMM, with the shaded area indicating the confidence interval. **B**, Group-level consistent d-PAC networks (top 20% of connections) for preterm (33– 37 w), term (37–41 w), and post-term (41–45 w) age groups. Nodes are labelled according to their assignment to the sensorimotor (S, orange) or association (A, magenta) regions. The directionality of d-PAC is visualised by arrows, with the tail denoting the region providing the low-frequency component and the arrowhead indicating the region providing the high-frequency component. For clarity, arrowheads are coloured according to the regional assignment of the low-frequency node. The diagrams below the glass brains summarise connection densities within and between S and A regions. **C**, Distributions of connection lengths for the age-specific consistent networks and the full network (grey; comprising all parcel-parcel connections). The distribution for each age group was compared with that of the full network using K-S tests, where D represents the K-S test statistic, and P values are shown after Benjamini-Hochberg correction. These results correspond to the theta band during QS; all other band-state combinations are shown in Supplementary Fig. 2.

### Global burst–framed connectivity declines despite greater sensorimotor– association connectivity after term

The global levels of burst-framed connectivity (NPPC; middle panel in Fig. 1C) showed a robust developmental decline (R^2^ = 0.42; P < 0.001; Fig. 3A) throughout the studied age span. However, spatial organization of these networks showed regionally selective development. While the local, within-region network densities peaked around term age (4.1% for within-S and 2.1% for within-A; Fig. 3B), there was a steady developmental increase in the connection density between A and S regions (1.7% vs 1.8% vs 2.3%; see Fig. 3B). This change towards denser S-A connectivity is also salient in the network topologies (Fig. 3B). Moreover, a clear developmental change was seen in the length distribution of burst-framed connectivity where early preference of short connections (with a peak at 3-4 cm; Fig. 3C) was gradually replaced by an increasing proportion of mid- and long-range connections (Fig. 3C). All these results were essentially consistent across alternative thresholds, frequency bands and sleep states (Supplementary Figs. 1B and 3). Taken together, burst-framed connectivity showed a robust global decline with increasing long-range connectivity between sensorimotor and association regions.

**Figure 3.**
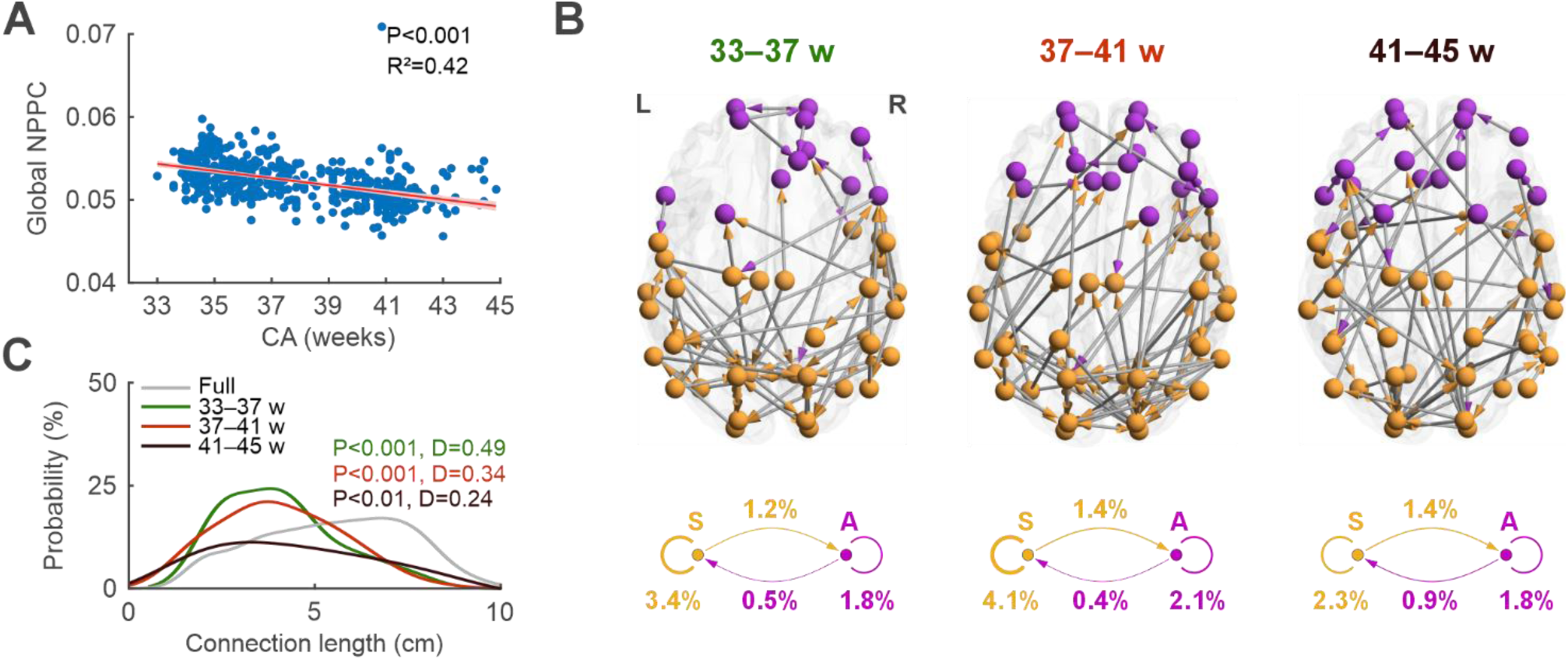
Development of burst-framed connectivity (NPPC). **A**, Developmental decline in global NPPC strength. Individual points correspond to EEG recordings, while the red line depicts the fitted LMM, with the shaded area indicating the confidence interval. **B**, Group-level consistent NPPC networks (top 20% of connections) for preterm (33–37 w), term (37–41 w), and post-term (41–45 w) age groups. Nodes are labelled according to their assignment to the sensorimotor (S, orange) or association (A, magenta) regions. The analytic asymmetry of NPPC is indicated by arrows, with the tail denoting the node at which both the low- and high-frequency components were considered (see Fig.1C). The diagrams below the glass brains summarize connection densities within and between S and A regions. **C,** Distributions of connection lengths for age-specific consistent networks and the full network (grey; comprising all connections). The distribution for each age group was compared with that of the full network with K-S tests, where D denotes the K-S test statistic, and P values are shown after Benjamini-Hochberg correction. The results are shown for the theta band during QS. For other frequencies and sleep states, see Supplementary Fig. 3.

### Phase–based connectivity shows global decrease in strength with consistent hierarchical organization

Similar to burst-related measures above, the global levels of phase-based connectivity (PPC; bottom panel in Fig. 1C) decreased with age (R^2^ = 0.56; P < 0.001; Fig. 4A). While the overall spatial distribution of consistent PPC networks was essentially unchanged over development, the strongest and weakest connections exhibited clearly diverging density and length profiles (Fig. 4A-C): The strongest connections were predominantly short-range, and substantially denser within the A and S regions (3.3–4.4%) than between them (0.7–1.2%; Fig. 4B). By contrast, the weakest connections were predominantly long-range, and substantially sparser within the A and S regions (0.5–1.6%) than between them (7–8.6%; Fig. 4B). The spatial distribution of participating nodes was also broadly similar across age groups. The PPC findings were robust to different thresholds and qualitatively consistent across all frequency bands and sleep states (Supplementary Figs. 1C, 4, and 5). As a technical verification, changes in PPC levels were not confounded by developmentally changing signal amplitudes (4–22 Hz: rho < 0.12; Supplementary Fig. 6).

**Figure 4.**
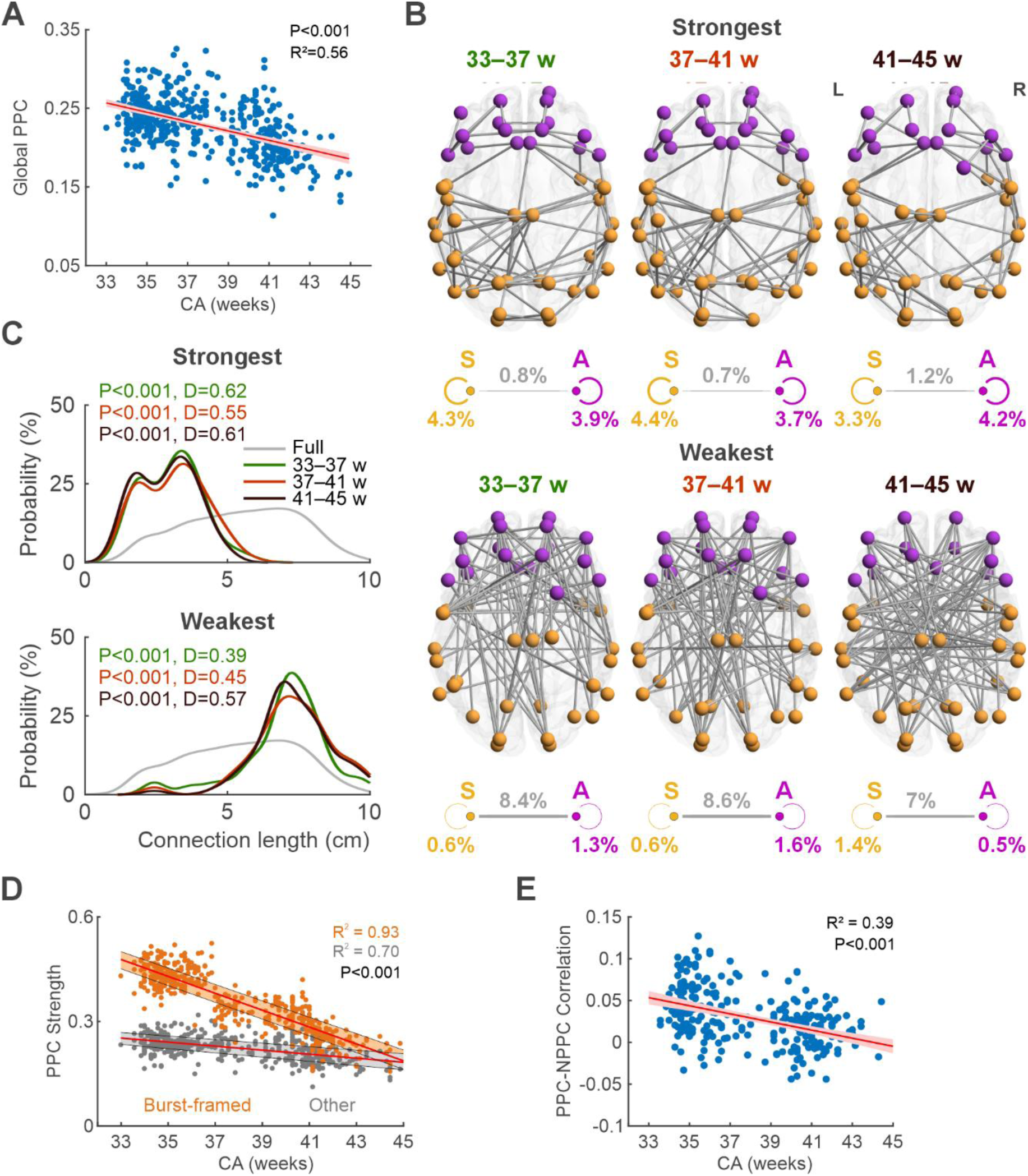
Development of phase-based connectivity (PPC). **A**, Developmental decline in global PPC strength. Each point denotes the mean PPC of one EEG recording, and the red line depicts the fitted LMM, with the shaded area indicating the confidence interval. **B**, PPC consistent networks for preterm (33–37 w), term (37–41 w), and post-term (41–45 w) age groups. Nodes are labelled according to their assignment to sensorimotor (S, orange) or association (A, magenta) regions. The top row shows group-level connections consistently ranked among the strongest 20% across individuals, while the bottom row shows those ranked among the weakest 20%. The diagrams below the networks summarize connection densities within and between S and A regions. **C**, Distributions of connection lengths for the strongest and weakest age-specific consistent networks and the full network (grey). The distribution for each age group was compared with that of the full network using K-S tests, where D denotes the K-S test statistic and P the Benjamini-Hochberg-corrected p-value. **D**, Developmental trajectories of mean PPC strength for connections belonging to the NPPC consistent networks (orange; ‘Burst-framed’; see also Fig.3B) compared with all other connections (grey; ‘Other’). The red lines depict the fitted LMMs, and the shaded areas indicate 25th–75th percentiles for each group. **E**, Change in PPC-NPPC correlation with age. For each EEG recording, the individual PPC-NPPC correlation (Spearman) was computed across all connections. The red line depicts the fitted LMM, with the shaded area indicating the confidence interval. Results are shown for the theta band during QS; all other frequency bands and states are presented in Supplementary Figs. 4 and 5.

### Phase-based connectivity develops increasing burst-independence around term age

Finally, we assessed our core hypothesis that cortical bursts provide the spatio-temporal frames for training the brain-wide PPC networks. Due to the global decline in all the absolute connectivity levels (see above), this hypothesis needs to be assessed within spatial contexts. Analytically, this would correspond to a developmental passage where PPC in the early networks show spatial and temporal correlation to burst occurrence (high NPPC), whereas later PPC is decreasingly correlated to the NPPC levels of the given connections. The findings indicate a clear developmental difference in the PPC strengths within vs outside of the burst-framed NPPC networks (see Fig. 3A). In early preterm infants, PPC is initially much stronger in burst-framed networks and showed a robust developmental decline until the PPC strengths became comparable within and between bursts after term age (regression estimates crossing at ∼45 weeks; Fig 5A). A complementary, connection-wise analysis across all possible network connections in the whole study cohort (3306 connections) showed a significant age-related decrease in PPC-NPPC correlations (R^2^ = 0.39; P < 0.001; Fig. 4D), indicating declining spatial and temporal dependence of PPC connectivity on the cortical bursting activity.

### Burst-framed phase synchrony associates with the later developing higher-order brain functions

The findings above showed that cortical bursts provide spatial and temporal frames for emerging phase-based connectivity (PPC), which is indicated in higher-order brain functions (Grover et al., 2021; Kohn et al., 2020). Next, we assessed whether the PPC strengths within the burst-framed connections would link to long-term neurocognitive development assessed using Bayley Scales at 18 months (Fig. 5). We found a significant correlation of burst-framed PPC to both cognitive scores (QS: rho = −0.45, P = 0.005, P_FDR_ = 0.015; AS: rho = −0.35, P = 0.03, P_FDR_ = 0.09) and language scores (rho = −0.49, P = 0.02, P_FDR_ = 0.03), while the composite motor performance showed no significant associations (Fig. 5). See Supplementary Fig. 7 for findings at other frequencies. These findings were specific to burst-framed connections, since the global mean PPC for the theta band in both sleep states was not associated with neurodevelopmental scores (P > 0.12 for all) suggesting that the observed associations were specific to burst-framed connections.

**Figure 5.**
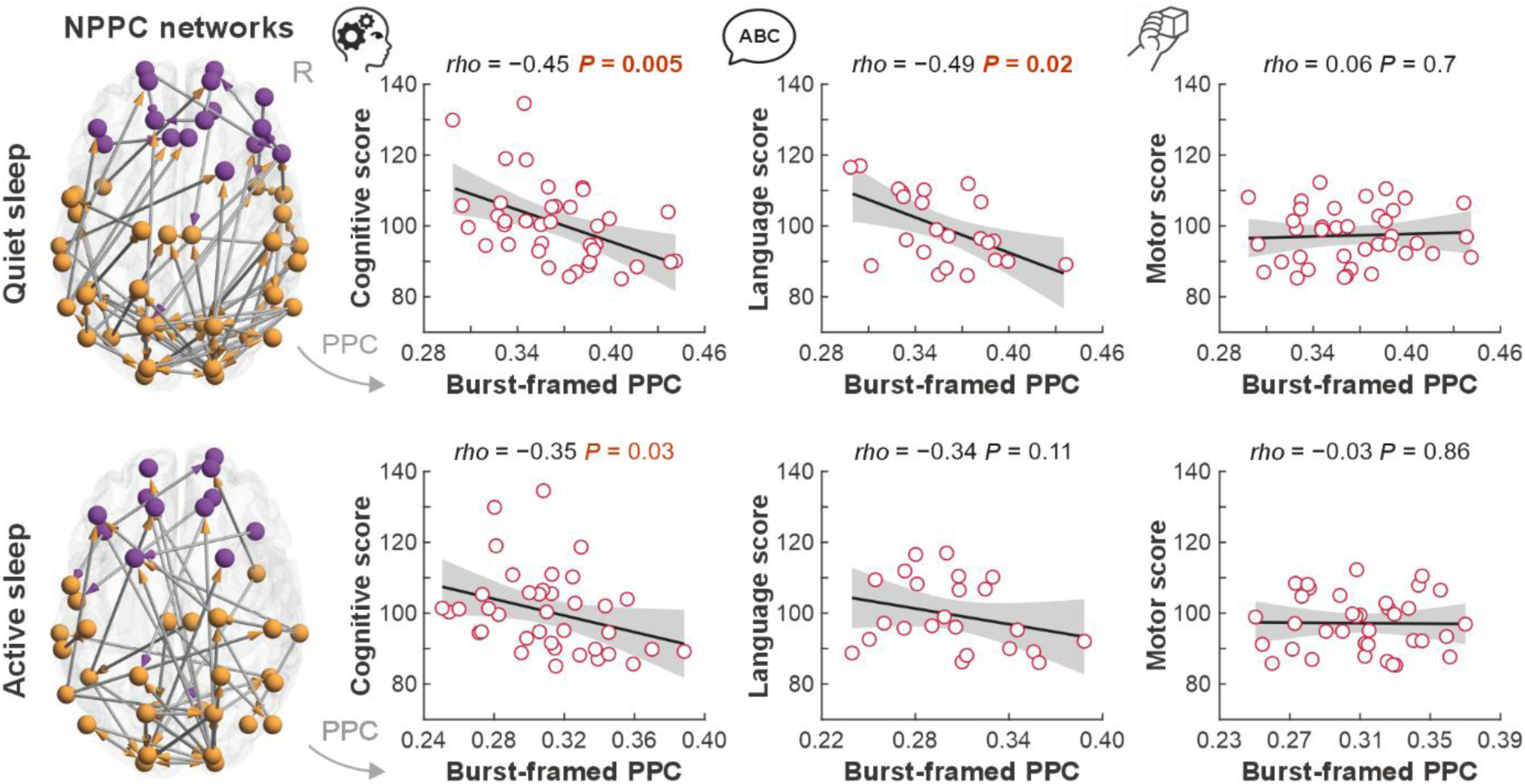
Association between burst-framed phase synchrony and later neurodevelopment. Glass brains show the NPPC consistent networks in the theta band (4–8 Hz) around term age (37–41 weeks). Nodes are labelled according to their assignment to association (A, magenta) or sensorimotor (S, orange) regions. Each point in the scatter plots represents an individual infant, showing mean PPC strength along the corresponding burst-framed connections against the respective Bayley score. The black lines show the fitted linear models, with the shaded areas indicating the 95% confidence intervals. Associations were assessed using Spearman correlations. Uncorrected significant P values are shown in red, whereas those remaining significant after Benjamini-Hochberg correction are shown in bold red.

## Discussion

Our findings reveal robust developmental changes in large-scale cortical network mechanisms around the birth transition: a shift from initially burst-based to later burst-framed connectivity, followed by increasing burst-independence in phase synchrony, the most temporally precise mode of corticocortical connectivity. These mode-specific network changes differed along the hierarchical S-A axis, and phase synchrony along burst-framed connections was associated with later neurodevelopmental outcomes. Here, we extend prior work by testing a mechanistic hypothesis that links the well-established cortical activity bursts to the gradual, spatio-temporally selective refinement of precise phase-based neural connectivity, a presumed underpinning of many higher-order brain functions (Fries, 2015; Grover et al., 2021; Kohn et al., 2020).

Since better neurocognitive performance is generally associated with stronger phase synchrony (Panda et al., 2021; Sato et al., 2018; Violante et al., 2017), it may seem counter-intuitive that early brain development is characterized by a robust global decline in all three connectivity measures. However, our findings are in line with prior reports spanning fetal and neonatal brain development (van de Pol et al., 2018; Varley et al., 2025; Yrjölä et al., 2024), and may arise from both biophysical changes and increasing spatio-temporal specificity of network organization (Chen et al., 2024; Larivière et al., 2020; Wu et al., 2024). The global strength of EEG-derived connectivity measures may partly reflect developmental changes in cortical geometry and the conductive pathways of cranial tissues (Azizollahi et al., 2016; Odabaee et al., 2014). At the same time, maturation of inhibitory circuitry (Chini et al., 2022; Wu et al., 2024) may increasingly constrain widespread cortical activity and facilitate more selective network interactions. More generally, the developmental decline in global connectivity may accompany increasingly selective organization of connections across the cortical hierarchy, en route to supporting the later emergence of higher-order brain functions (Gao et al., 2015; Li et al., 2024; Luo et al., 2024).

Importantly, the spatial organization of corticocortical connectivity developed differently across the three modes. Burst-based connectivity (d-PAC) shifted from a widespread, long-range pattern towards more spatially confined and segregated connectivity within S and A regions. In contrast, burst-framed connectivity (NPPC) shifted from initially strong within-S networks towards increasing connectivity along the S-A axis, consistent with the gradual transition from predominantly sensory to more multimodal cortical networks (Li et al., 2024; Nava et al., 2024). PPC showed comparatively stable spatial organization with respect to the S-A hierarchy. This stability, with the strongest connections preferentially linking regions within the same broad cortical division and the weakest connections more often spanning the S-A axis, suggests that general phase synchrony already exhibits a well-organized spatial topology during this developmental period. These distinct spatial trajectories suggest that maturation does not simply strengthen or weaken cortical connectivity, but involves mode-specific reorganization of cortical networks.

Experimental literature provides support and possible mechanistic explanations for these observations. Early cortical bursting is predominantly found in sensorimotor cortices and is driven by sensory inputs (Gómez et al., 2021; Luhmann et al., 2016; Martini et al., 2021), and guides the burst-based, activity-dependent maturation in corticocortical networks (Molnár et al., 2020; Mukherjee and Kanold, 2022). One possible interpretation is that the early emergence of NPPC in sensorimotor regions reflects the strong contribution of immature thalamocortical inputs engaging transient subplate circuitry (Borden et al., 2022; Molnár and Kwan, 2024). This circuitry integrates endogenous and sensory-driven activity and supports early cortical networks through the burst-like activity patterns observed in preterm EEG (Khazipov and Luhmann, 2006; Kostović and Judas, 2010; Molnár et al., 2020; Tolner et al., 2012; Vanhatalo and Kaila, 2006). Specifically, early activity bursts driven by common thalamic input to the subplate may provide temporally precise and spatially constrained references for high-frequency synchrony within sensory-dominant networks (Kostka et al., 2020), whereas early long-range d-PAC between association and sensorimotor regions may reflect a transient scaffold coordinating activity between these regions (Kostović and Judas, 2010; Molnár et al., 2020).

With maturation, the shift of d-PAC towards more local connectivity within S and A regions, together with the reorganization of NPPC towards greater connectivity along the S-A axis and greater involvement of A regions after term age, is consistent with both the decline of transient burst-centred communication and the rapid reorganization of neonatal functional networks across the birth transition (Burlando et al., 2026; Riyahi et al., 2021; Tokariev et al., 2016b; Yrjölä et al., 2024). These findings are also in line with structural work showing that basic thalamocortical organization is established early, while perinatal thalamocortical and longer-range corticocortical connectivity continues to mature across this developmental window (Keunen et al., 2017; Kostović et al., 2021; Oldham et al., 2025).

Perhaps the most striking observation is the coupling of PPC to burst-framed (NPPC) networks spanning the evolutionarily conserved hierarchical S-A axis. PPC was preferentially concentrated along burst-framed connections early in development, and this relationship weakened with maturation. These findings are compatible with a spatially and temporally specific role for cortical bursts in the development of corticocortical phase synchrony. Specifically, burst-framed activity may initially provide a temporal framework for phase synchrony along selected cortical connections before PPC becomes progressively more burst-independent with maturation.

The functional relevance of these developmental network dynamics is further supported by their association with later neurodevelopmental outcomes. Stronger PPC along burst-framed connections around term age was significantly associated with poorer later cognitive and language scores. Importantly, these associations were not observed for global PPC, suggesting that spatially specific phase synchrony along burst-framed connections may provide a more informative marker of cortical network maturation. Given that PPC became progressively less coupled to burst-framed organization with age, greater persistence of phase synchrony along burst-framed connections around term age may reflect a less mature network state or a delayed transition towards burst-independent synchrony. These findings are in line with previous studies linking neonatal functional connectivity to later neurobehavioral outcomes (Shellhaas et al., 2022; Toulmin et al., 2021; Yrjölä et al., 2021).

Several limitations should be considered when interpreting these findings. First, the observed associations between cortical bursting, phase synchrony and later neurodevelopment cannot establish causal relationships or their underlying mechanisms. Second, the inherent spatial uncertainty of EEG source reconstruction and the assignment of cortical parcels to S and A regions introduce unavoidable ambiguity near regional boundaries. (Azizollahi et al., 2016). Third, phase synchrony was quantified using PLV to retain near-zero-lag interactions and to maintain methodological consistency with the other metrics, but it may remain susceptible to volume-conduction-related source leakage, particularly for the shortest-range connections (Palva and Palva, 2012). Fourth, all infants were born preterm, which allowed us to study development across the period spanning the typical birth transition; however, prematurity itself may affect connectivity mechanisms (Tokariev et al., 2019), although overall EEG development is known to follow postmenstrual age more closely than postnatal age (Stevenson et al., 2020).

In conclusion, our findings are consistent with the hypothesis that cortical bursting contributes to the emergence of phase-based synchrony in the developing preterm brain. The results further suggest that this process is accompanied by increasing spatial specificity and progressive independence of phase synchrony from burst-related activity with maturation. Together, these findings provide a framework for understanding how transient cortical bursting may scaffold the maturation of large-scale functional networks.

## Methods

### Subjects

Serial recordings of high-density EEG data were collected from sleeping preterm infants at 33–45 weeks of conceptional age (CA; term age is ≥ 37 weeks) to study early development of cortical modes in neuronal connectivity. The study dataset was collated from two clinical trials (ClinicalTrials.gov NCT01439269 (Welch et al., 2014), ClinicalTrials.gov NCT02710474 (Welch et al., 2022)); using identical recording setups across 496 recordings from 233 infants (gestational age at birth: 30.75 ± 2.23 weeks, mean ± STD). See Supplementary Fig. 8 for further details about the subjects.

### EEG Recordings

The EEGs were recorded using a 128-lead Electrical Geodesics EEG system (EGI Inc., Eugene, Oregon) in the intensive care unit during daytime sleep (∼60min), including at least one full sleep cycle with both active and quiet sleep states. Electrode drying was reduced with a plastic wrap, and the electrode impedances were maintained below 50 kΩ (Welch et al., 2014). The recording reference was placed at vertex, sampling rate was Fs = 1000 Hz, and band-pass filtering was from 0.1–400 Hz. All EEGs were converted to average montage and exported to European Data Format (EDF) for further processing.

### EEG pre-processing

All EEG recordings were first reviewed visually by trained experts (PY and SV) to identify good-quality epochs from periods of active sleep (AS; precursor of rapid eye movement sleep) and quiet sleep (QS; precursor of non-rapid eye movement sleep) states as described before (Yrjölä et al., 2026, 2024). Five minutes of clean data for each recording for both AS and QS (444 and 462, respectively) were selected. Further EEG processing was done on these selected AS and QS segments only. First the signal baseline offset was removed by subtracting its mean and the signal was then band-pass filtered at 0.4–40 Hz using a pair of fifth-order Butterworth filters in both forward and backward directions to avoid phase shifts. Next, the EEG signals were down-sampled to Fs = 100 Hz. From all recordings four facial electrodes were excluded leaving 124 EEG channels per subject. Then, the signal quality of the data was visually re-inspected for notable artifacts (loose electrode contact, electromyogram (EMG), movements, other excessively high-amplitude), and poor channels and/or time windows were discarded. Finally, the electrocardiographic (ECG) artifacts were removed with a previously developed, semi-automated approach based on independent component analysis (ICA) (Hamaneh et al., 2014; Yrjölä et al., 2022). In some subjects, ICA was also used to remove electrical interference artifacts.

### Reconstruction of cortical signals

The EEG sensor signals were transformed to cortical-level source space signals using a three-compartment (scalp, skull, and intracranial volume) infant boundary element method (BEM) head model (Tokariev et al., 2019). To minimize model-dependent influence on the studied developmental change in cortical activity, we opted to use full-term infant cortex as the common cortical source template across all the datasets. We also cross-validated robustness of our key findings using a head model with a smoother cortical curvature that corresponds to approximately 35 weeks of gestation (Supplementary Fig. 9). The cortical source space was represented by 8014 dipoles oriented orthogonally to the surface, and all other shells were approximated by 2562 vertices each. The following conductivity values were used for infant head tissues: scalp 0.43 S/m, skull 0.2 S/m, and intracranial volume 1.79 S/m (Despotovic et al., 2013; Odabaee et al., 2014; Tokariev et al., 2016a). We computed the forward operator using symmetric boundary element method as implemented in the OpenMEEG package (Gramfort et al., 2010; Kybic et al., 2005). The inverse operator was then computed with dynamic statistical parametric mapping (Dale et al., 2000) as it is implemented in Brainstorm software (Tadel et al., 2011). Cortical sources were then clustered into 58 parcels according to the infant parcellation scheme (Tokariev et al., 2019b), which were further assigned to sensorimotor (S) and association (A) regions (Fig.1E) (Kolb, 2025; Larsen et al., 2023; Stevenson et al., 2025; Sydnor et al., 2023). Corresponding parcel signals were computed as a weighted mean of source signals within them (Tokariev et al., 2019). Finally, we divided all infants into three groups according to the age of the subjects at EEG recording: preterm (33–37 weeks, N = 192/191 at AS/QS), term (37–41 weeks, N = 127/141 at AS/QS), and post-term (41–45 weeks, N = 71/78 at AS/QS). To account for the non-independence of repeated measures, we averaged the corresponding metrics for subjects with multiple measurements within the same age group.

### Distant phase-amplitude coupling (d-PAC)

Burst-based cross-frequency connectivity, d-PAC, was analyzed between cortical signal pairs using a modified measure of phase-amplitude coupling (PAC; see Fig 1A; (Tokariev et al., 2016b; Vanhatalo et al., 2005) that assessed synchronization between phase of the low frequency wave in one cortical signal and the amplitude of higher frequency activity in another cortical signal. In other words, how bursts of higher frequency activity are “nested” within low frequency “nesting” oscillations. To do so, all parcel signals were filtered into slow ‘nesting’ (0.4–1.5 Hz) and fast ‘nested’ (4–8 Hz, theta; 8–13 Hz, alpha; 13–22 Hz, beta) oscillatory components using pairs of Butterworth filters (stopband attenuation at least 15 dB in one direction). Hilbert transform was used to obtain the high frequency activity amplitude envelopes, which were then filtered using the slow component setting (0.4–1.5 Hz). Finally, phase locking values (PLV) (Lachaux et al., 1999) were estimated between the phase of the slow component (in one cortical region) and the phase of the fast component amplitude envelope (in another cortical region), yielding d-PAC that reflects “burst-based” corticocortical connectivity (see Fig 1C). The resulting d-PAC connectivity matrices are asymmetric and visualized as directed networks.

### Nested phase-phase correlation (NPPC)

To test our hypothesis that low-frequency burst-based connectivity precedes and perhaps entrains more temporally precise phase synchronization (Canolty and Knight, 2010; Hyafil et al., 2015; Yrjölä et al., 2024), we developed a novel measure, nested phase-phase correlation (NPPC), to capture a putative intermediate burst-framed mode between d-PAC and PPC (Fig. 1B). For each pair of cortical signals, the slow nesting component (0.4–1.5 Hz) was computed from one cortical signal and the fast nested components were computed from both cortical signals (for example, 4–8 Hz). Then, phase-phase correlation (PPC) was computed with PLV (Lachaux et al., 1999) between the faster signal components, using 1-sec sliding windows and 0.1 second step size as a compromise between number of samples per window, temporal resolution, and computational speed. The PLV(t) time series were up-sampled to the original sampling rate (Fs = 100 Hz) and band-pass filtered using the same filters as for the slow nesting component (0.4–1.5 Hz). Finally, NPPC was computed as PLV between the slow nesting component and PLV(t). The outcome is an asymmetric, directed NPPC matrix where cortical parcel with the slow component is defined as the origin of each connection (arrows in the network plots were visualized accordingly). Taken together, this analytical process yields a measure of how phase synchrony in higher-frequency cortical activity is temporally aligned with the timing of bursts reflected in the lower-frequency component. Matlab code for computing NPPC is provided as part of the full analytical pipeline (see Code availability).

### Phase-phase correlation (PPC) networks

The phase-phase correlation (PPC) was estimated between cortical signal pairs as previously described (Tokariev et al., 2016b). In brief, phase difference (ΔΦ) was computed for each pair of corresponding phase time series, obtained with Hilbert transform from the band-pass filtered signals, and PPC was estimated using classical phase-locking value (PLV, (Lachaux et al., 1999)), defined as:

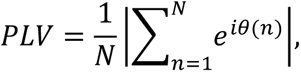

We chose PLV to support methodological consistency across our measures, as PLV is central for both d-PAC and NPPC. Due to the possibility of spatial leakage in PLV applied to source space signals (Palva and Palva, 2012), we cross-validated our PLV-based results by comparing developmental changes in global PPC with corresponding results obtained using debiased weighted phase lag index (Vinck et al., 2011) in our previous work. (Yrjölä et al., 2024). Further validation of the PLV-based PPC measures included assessing its amplitude-sensitivity using randomly selected data subset (Supplementary Fig. 6). Nevertheless, we interpret the detailed findings cautiously and considering the corticocortical distances.

### Computation of the network ‘backbone’

Consistent networks were defined as spatially consistent subsets of the strongest or weakest connections across subjects, representing the group-level network ‘backbone’ (Tokariev et al., 2019). For each subject, we first binarized the connectivity matrix by taking the top k% strongest, or the bottom k% weakest connections, depending on the analysis. We then identified connections that occurred more consistently across subjects than expected by chance. For each connection, we summed up the individual binary matrices across subjects and tested whether the observed number of subjects showing that connection exceeded chance expectation using binomial statistics (significance threshold α = 0.05). The resulting significant connections were further controlled for multiple comparisons by rejecting the fraction of connections with the highest p-values (Palva et al., 2010). This fraction was set to 5% of the total number of tested connections, corresponding to the rejection of up to N = 83 connections for PPC networks, which were represented by symmetric matrices, and up to N = 166 connections for d-PAC and NPPC, which were represented by asymmetric matrices. We repeated the analysis using thresholds of k = 10%, 20%, and 30% and found that the resulting network patterns were robust across thresholds (Supplementary Fig. 1). For all resulting networks, we quantified the density of connections within and between parcels assigned to association and sensorimotor cortices. Density was calculated as the percentage of backbone connections relative to the total number of possible connections within or between the corresponding cortical classes.

Finally, we also compared PPC strength within the NPPC networks with PPC strength in the rest of the cortex. Within-network connectivity was computed by masking individual PPC matrices with the group-level NPPC templates and then averaging the resulting values. To estimate PPC levels in the rest of the cortex, we randomly selected the same number of PPC connections as in the NPPC network, but from outside the template, and computed their mean for each infant. This procedure was repeated 100 times for each subject, and the mean across iterations was taken as the final estimate. PPC levels within and outside NPPC networks were compared using Kolmogorov–Smirnov test (K–S test). In addition, for the NPPC networks, we investigated the change in correlation between NPPC and PPC with age. For this, we computed the Spearman correlation between NPPC and PPC in all connections and fitted an LMM model with the resulting coefficients.

### Assessment of network topology across sensorimotor-association cortices

In addition to the global-level general measures, we studied the emerging network topology using the “hierarchical patterning” context, motivated by recent studies showing that developmental gradients across the sensorimotor-to-association (S–A) axis are seen at multiple levels, from genetic expression to in vivo neuroimaging (Sydnor et al., 2021), including neuronal activity (Stevenson et al., 2025). Therefore, the S–A axis provides a physiologically relevant framework to study the cascades of functional neurodevelopment, including maturation of cortical neuronal activity (Stevenson et al., 2025). We adopted the previously established S-A assignment in our cortical template (see Fig S3 in (Stevenson et al., 2025)), where cortical activity parcels were determined to belong to the sensorimotor (S) or association (A) areas (Ball et al., 2020; Larsen et al., 2023). Then, we quantified the network measures used here (Fig. 1C) separately for connections within S or A regions (within-S and within-A) and between S and A regions (S–A).

### Associations of neuronal activity networks with neurodevelopment

We computed correlations between the strengths of PPC connectivity within the NPPC consistent networks (at term age; 37-41 weeks of CA) and neurodevelopmental outcomes using the well-established, standardized Bayley Scales scores (BSID-III) (Bayley, 2006) at 18 months. Following the common approach in studying typically developing children, we excluded scores of individuals below -1SD (cutoff of 85 points (Johnson et al., 2014)), yielding the following final sample sizes for subjects with both EEG data and corresponding Bayley scores: N = 38 for cognitive (in both states), N = 23 for language (in both states), and N = 36/38 (in AS/QS) for motor. The network-Bayley correlations were computed for all networks using nested frequency bands: theta (4–8 Hz), alpha (8–13 Hz), and beta (13–22 Hz).

### Statistical analysis

Distributions of connection lengths (Euclidean lengths of network edges) were compared between consistent networks and the full network (all possible connections) using two-sample Kolmogorov-Smirnov test. Correlations of network strengths to neurodevelopment scores were estimated using non-parametric Spearman correlation due to the non-normal distributions. We used Kolmogorov–Smirnov tests (K-S tests) to assess whether the distributions of subject-level mean PPC differed between connections within the NPPC network and global PPC after excluding those network connections. Developmental changes in global PPC, d-PAC, and NPPC connectivity levels were assessed using linear mixed models (LMM) (Yrjölä et al., 2024). These regression models capture linear effects, while allowing non-independence of repeated measures, inconsistent assessments, and subject-specific differences in starting level and coefficients (McCormick et al., 2023). False discoveries due to multiple comparisons were corrected using Benjamini-Hochberg method (BH), and the nominal significance level was set to p<0.05. The effect sizes were estimated with adjusted R^2^ values, and the models were separately computed for each frequency band and both sleep states.

## Code availability

Custom preprocessing and analyses were performed in MATLAB R2022b (MathWorks, Natick, MA). The analytical pipeline for computing connectivity metrics, together with the infant head model and network visualization scripts, is available at https://github.com/babyEEG/infant-phase-connectivity. Source reconstruction was performed using Brainstorm (Tadel et al., 2011) with OpenMEEG (Gramfort et al., 2010; Kybic et al., 2005). Independent component analysis for artefact removal was performed using FastICA ((Hyvärinen, 1999), https://research.ics.aalto.fi/ica/fastica/) and Picard (Ablin et al., 2018).

## Data Availability

The EEG data analysed in this study are available upon reasonable request through a data-sharing agreement with Columbia University Medical Center (NY, USA).

## Acknowledgements

This work was supported by Emil Aaltonen Foundation (S.K.), Sigrid Jusélius Foundation (P.Y., S.V. and A.T.), Orion Research Foundation (P.Y.). We thank Prof. Martha G. Welch and Prof. Michael M. Myers from Columbia University Medical Center for providing the EEG and outcome data that were re-analysed in this work.

## Author contributions

Conceptualisation: S.V., A.T.; Data preprocessing: P.Y.; Data curation: S.K., P.Y.; Formal analysis: S.K., Funding acquisition: S.V., A.T.; Methodology: S.K., P.Y., S.V., A.T.; Software: S.K., P.Y., A.T; Supervision: A.T.; Validation: S.K.; Visualisation: S.K., S.V., A.T.; Writing original draft: S.K., S.V., A.T.; Reviewing and editing: P.Y.

## Supplement

**Supplementary Figure 1.**
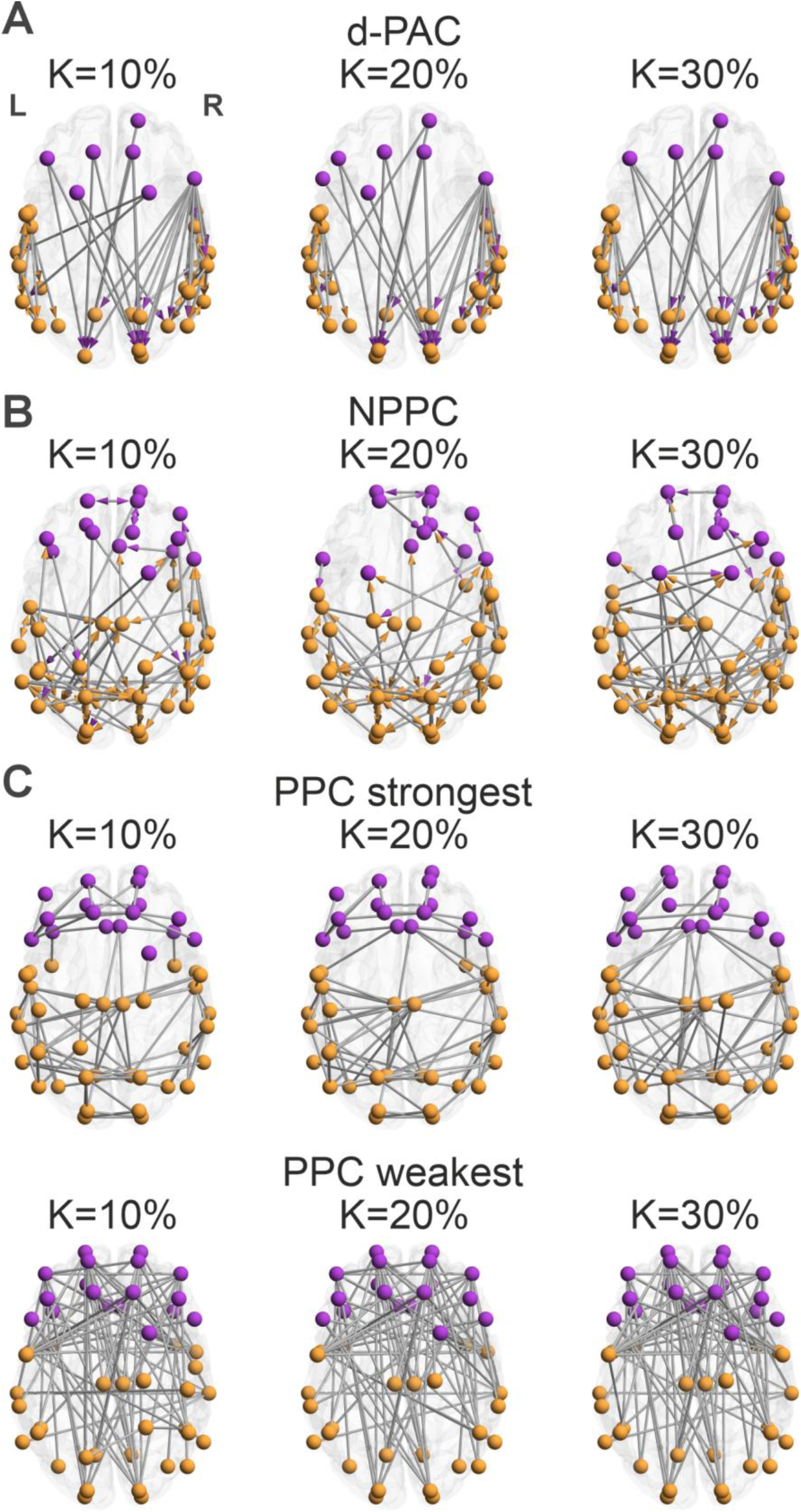
Consistent networks for theta band during QS in preterm (37–41 w) with varying thresholding (K=10%, 20%, and 30%) for d-PAC, NPPC, and PPC.

**Supplementary Figure 2.**
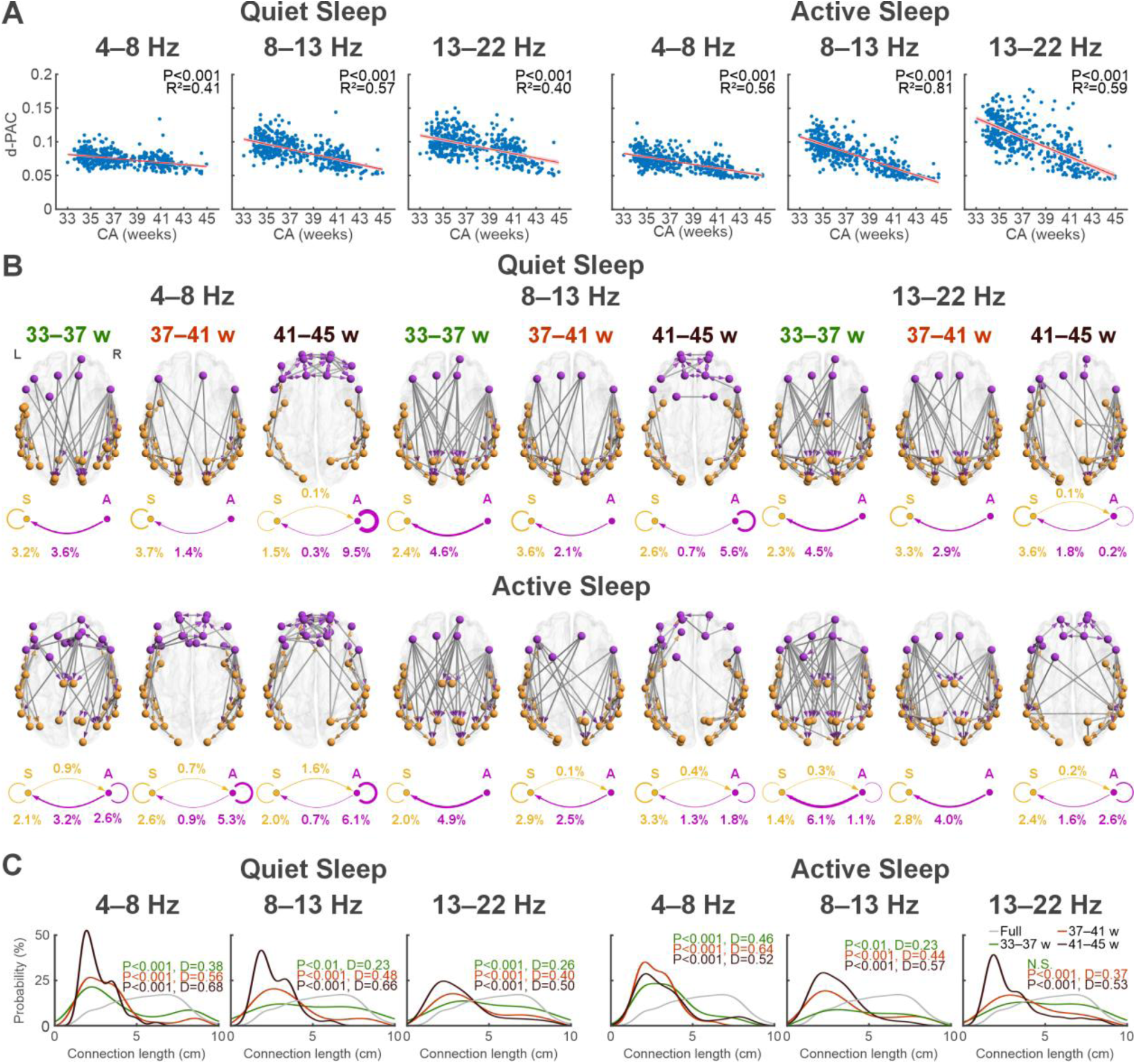
Development of burst-based connectivity. **A**, Depicts the developmental decline in global d-PAC levels, with each point depicting one EEG recording, and the red line depicting a fitted LMM model with the shaded areas indicating the confidence interval. **B**, Shows the consistent d-PAC networks (20% strongest) in the groups of preterm (33–37 w), term (37–41 w), and post-term age (41– 45 w) infants, with nodes labelled according to their assignment to the sensorimotor or association regions (S = orange, A = magenta). The asymmetry of d-PAC measure is reflected with the arrow base (low-frequency component) and the arrow point (high frequency component), respectively, while the arrow point color indicates the regional assignment (S vs A). The plots below the glass brains summarize connection densities between/within S and A regions, expressed as proportion of connections of all possible connections. **C,** Distribution of connection lengths of the consistent networks in the three age groups as well as the full network (grey; all parcel-parcel connections). All distributions were compared to the full network with K-S tests, where D represents the K-S test statistic, and P value is shown after Benjamini-Hochberg correction.

**Supplementary Figure 3.**
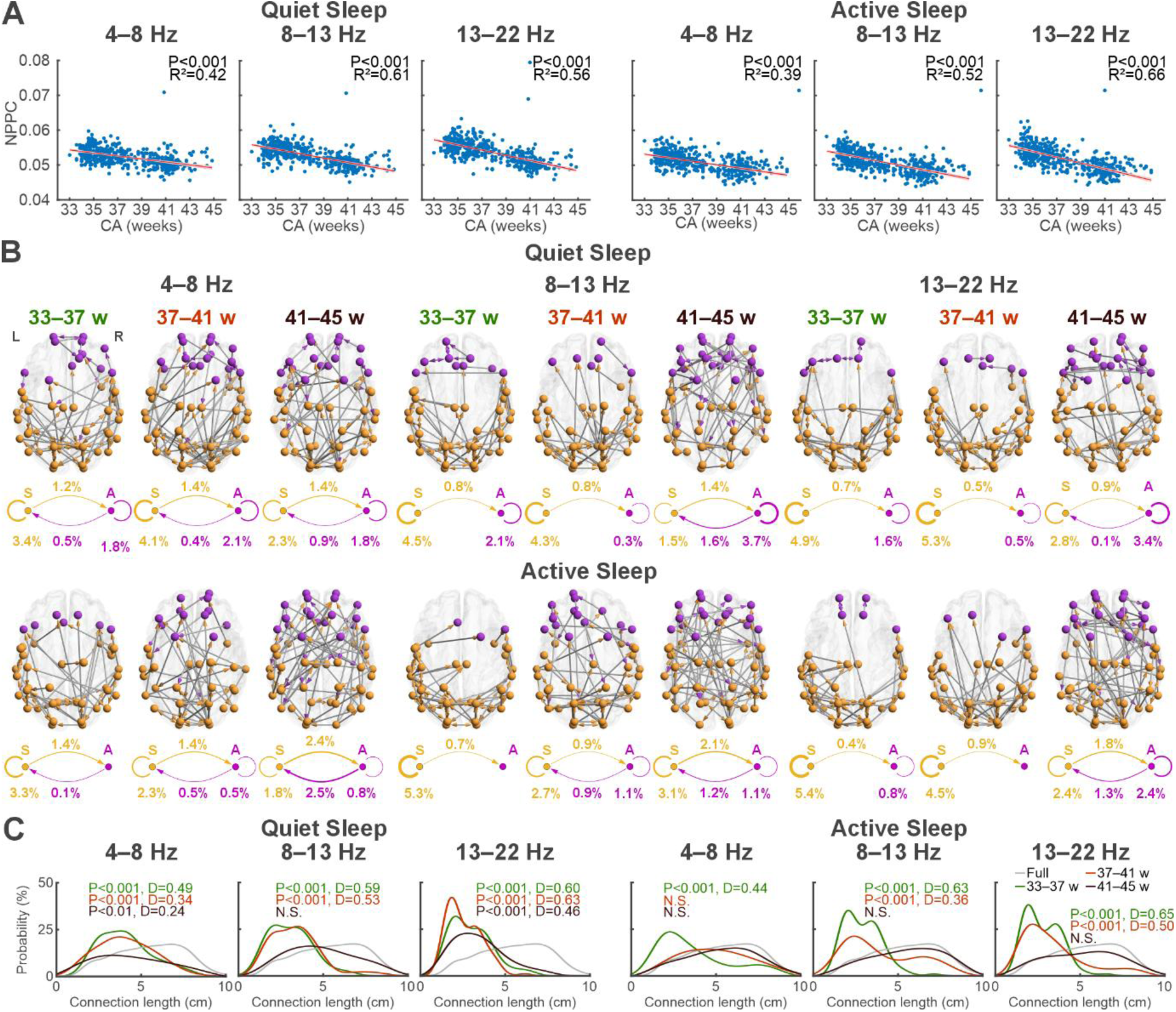
Development of NPPC consistent networks. **A**, Depicts the developmental decline in global NPPC levels, with each point depicting one EEG recording, and the red line depicting a fitted LMM model with the shaded areas indicating the confidence interval. **B**, Shows the consistent NPPC networks (20% strongest) in the groups of preterm (33–37 w), term (37–41 w), and post-term age (41– 45 w) infants, with nodes labelled according to their assignment to the sensorimotor or association regions (S = orange, A = magenta). The asymmetry of NPPC measure is reflected with the arrow, where the point of the arrows denotes the source without a nested low frequency component. The plots below the glass brains summarize connection densities between/within S and A regions, expressed as proportion of connections of all possible connections. **C,** Distribution of connection lengths of the consistent networks in the three age groups as well as the full network (grey; all parcel-parcel connections). All distributions were compared to the full network with K-S tests, where D represents the K-S test statistic, and P value is shown after Benjamini-Hochberg correction.

**Supplementary Figure 4.**
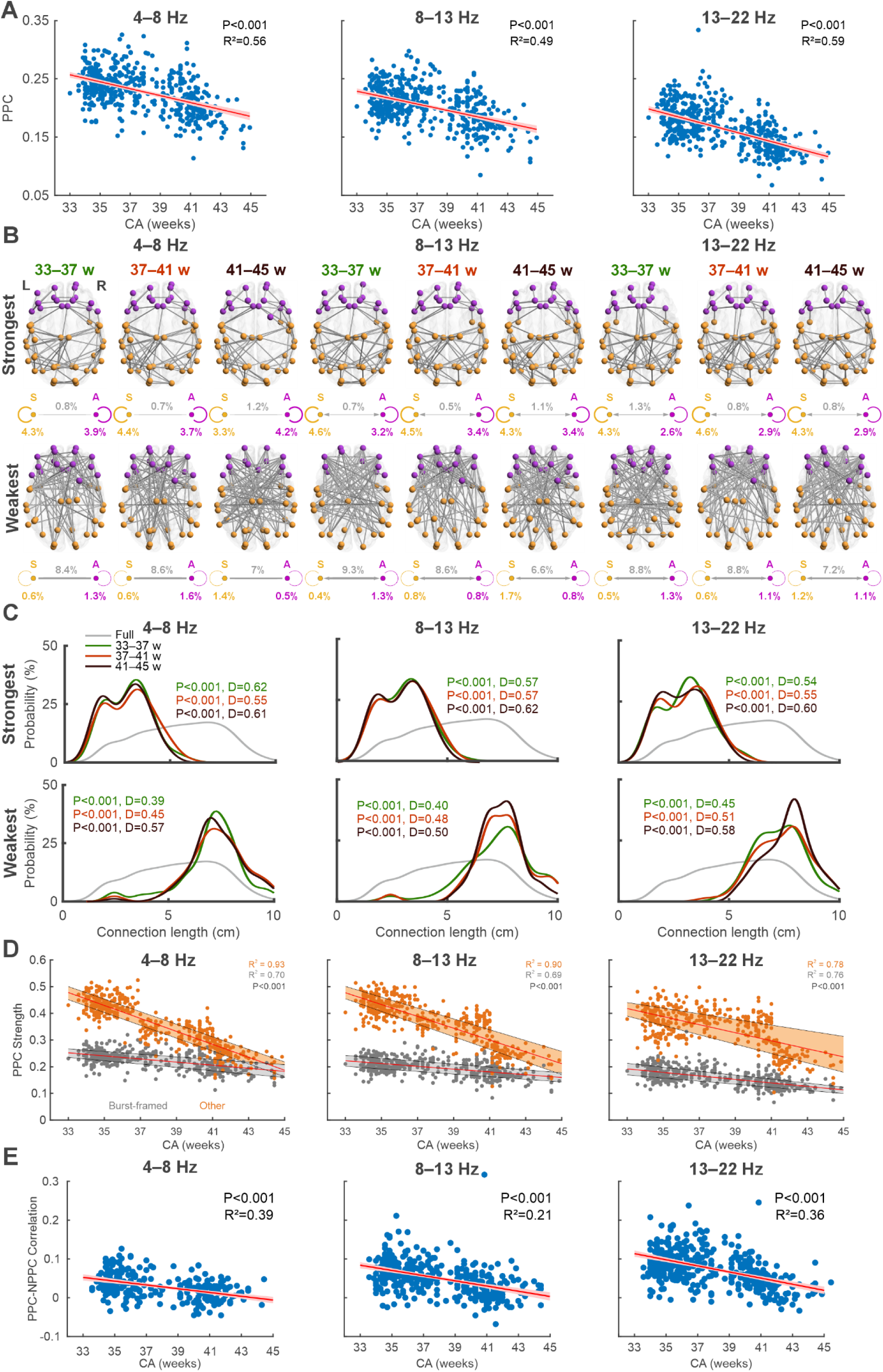
Development of PPC consistent networks during QS. **A**, The change in global PPC levels from preterm to post-term age (33–45 weeks CA). Each point denotes the mean PPC of one recording, and the red line depicts the fitted LMM model with the shaded areas indicating the confidence interval. **B**, PPC consistent networks for preterm (33–37 w), term (37–41 w), and post-term age (41– 45 w) infants. Nodes are labelled according to the S-A axis (S = orange, A = magenta). The top row shows the connections that are consistently in the top 20% strongest, and bottom row for the weakest 20%. Percentages below the networks display their density between/within sensorimotor (S, orange) and association (A, magenta) nodes. The density is computed as the proportion of connections between (S-A, grey) or within the S (S-S, yellow) and A (A-A, magenta) nodes of all possible connections between/within the regions. **C**, Distribution of connection lengths of the networks for all groups and the full network (grey). All distributions were compared to the full network with K-S tests, where P indicates the Benjamini-Hochberg-corrected p-value of the K-S test and D represents the test statistic of the K-S test. **D,** LMM models for mean PPC strength for connections belonging to the NPPC consistent networks (orange; ‘Burst-framed’; see also Fig.3B) compared with all other connections (grey; ‘Other’). The red line depicts the fitted LMM model and the shaded areas indicate 25–75 quantiles for each group. **E**, Change in PPC-NPPC correlation with age. For each EEG recording, the individual PPC-NPPC correlation (Spearman) was computed across all connections. The red line depicts the fitted LMM, with the shaded area indicating the confidence interval.

**Supplementary Figure 5.**
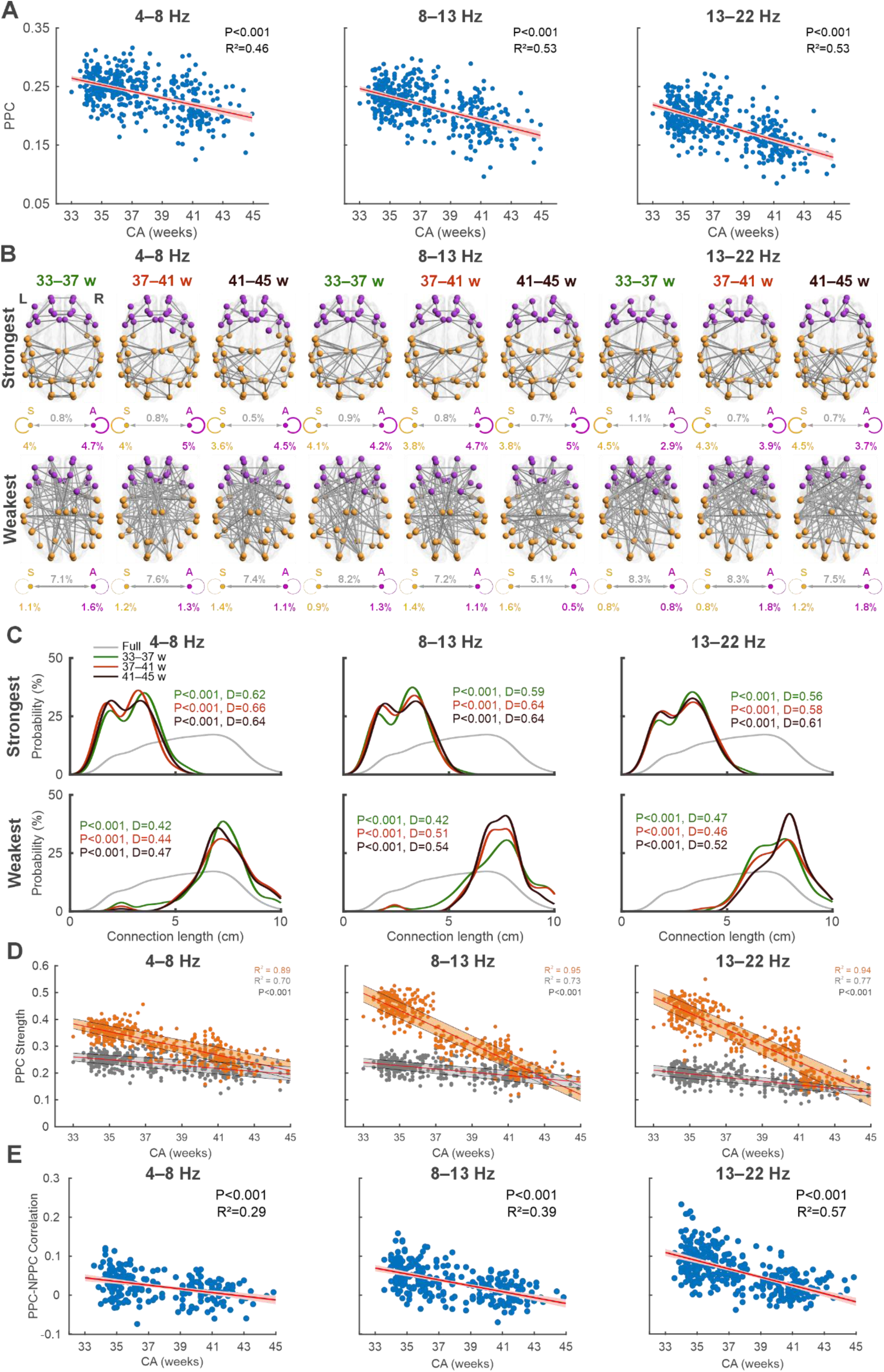
Development of PPC consistent networks AS. **A**, The change in global PPC levels from preterm to post-term age (33–45 weeks CA). Each point denotes the mean PPC of one recording, and the red line depicts the fitted LMM model with the shaded areas indicating the confidence interval. **B**, PPC consistent networks for preterm (33–37 w), term (37–41 w), and post-term age (41–45 w) infants. Nodes are labelled according to the S-A axis (S = orange, A = magenta). The top row shows the connections that are consistently in the top 20% strongest, and bottom row for the weakest 20%. Percentages below the networks display their density between/within sensorimotor (S, orange) and association (A, magenta) nodes. The density is computed as the proportion of connections between (S-A, grey) or within the S (S-S, yellow) and A (A-A, magenta) nodes of all possible connections between/within the regions. **C**, Distribution of connection lengths of the networks for all groups and the full network (grey). All distributions were compared to the full network with K-S tests, where P indicates the Benjamini-Hochberg-corrected p-value of the K-S test and D represents the test statistic of the K-S test. **D,** LMM models for mean PPC strength for connections belonging to the NPPC consistent networks (orange; ‘Burst-framed’; see also Fig.3B) compared with all other connections (grey; ‘Other’). The red line depicts the fitted LMM model and the shaded areas indicate 25–75 quantiles for each group. **E**, Change in PPC-NPPC correlation with age. For each EEG recording, the individual PPC-NPPC correlation (Spearman) was computed across all connections. The red line depicts the fitted LMM, with the shaded area indicating the confidence interval.

**Supplementary Figure 6.**
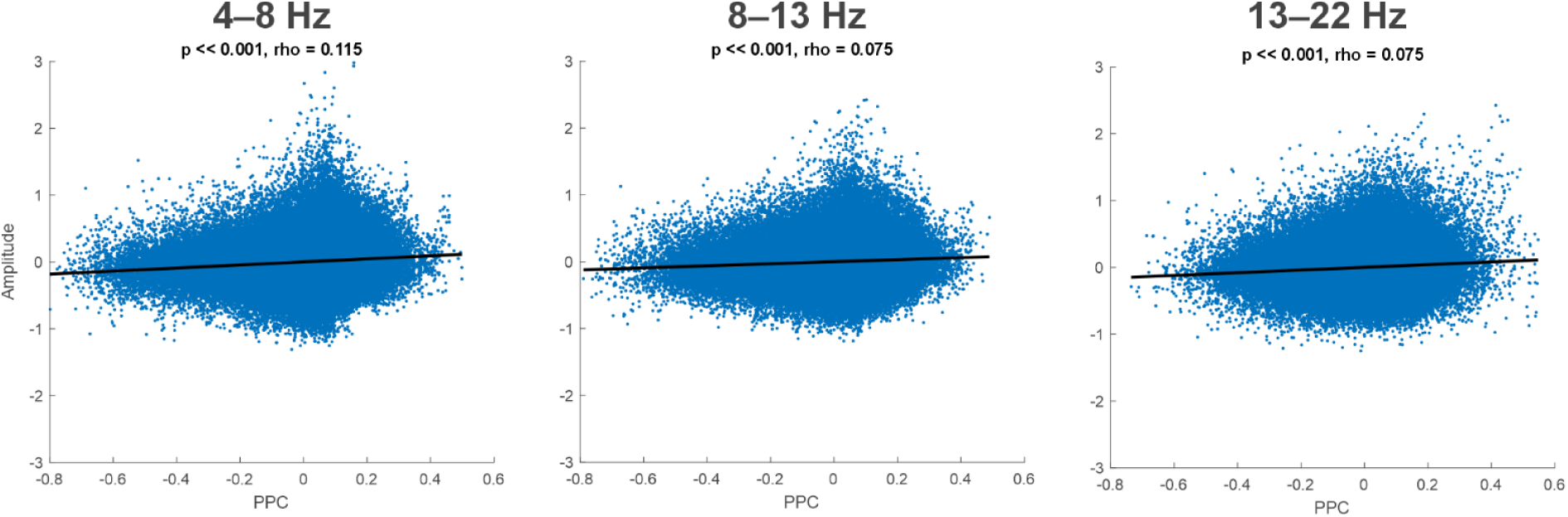
Correlation of PLV from ten subjects in ten connections in ten second windows for two minutes with the corresponding amplitude of the parcel signals during QS.

**Supplementary Figure 7.**
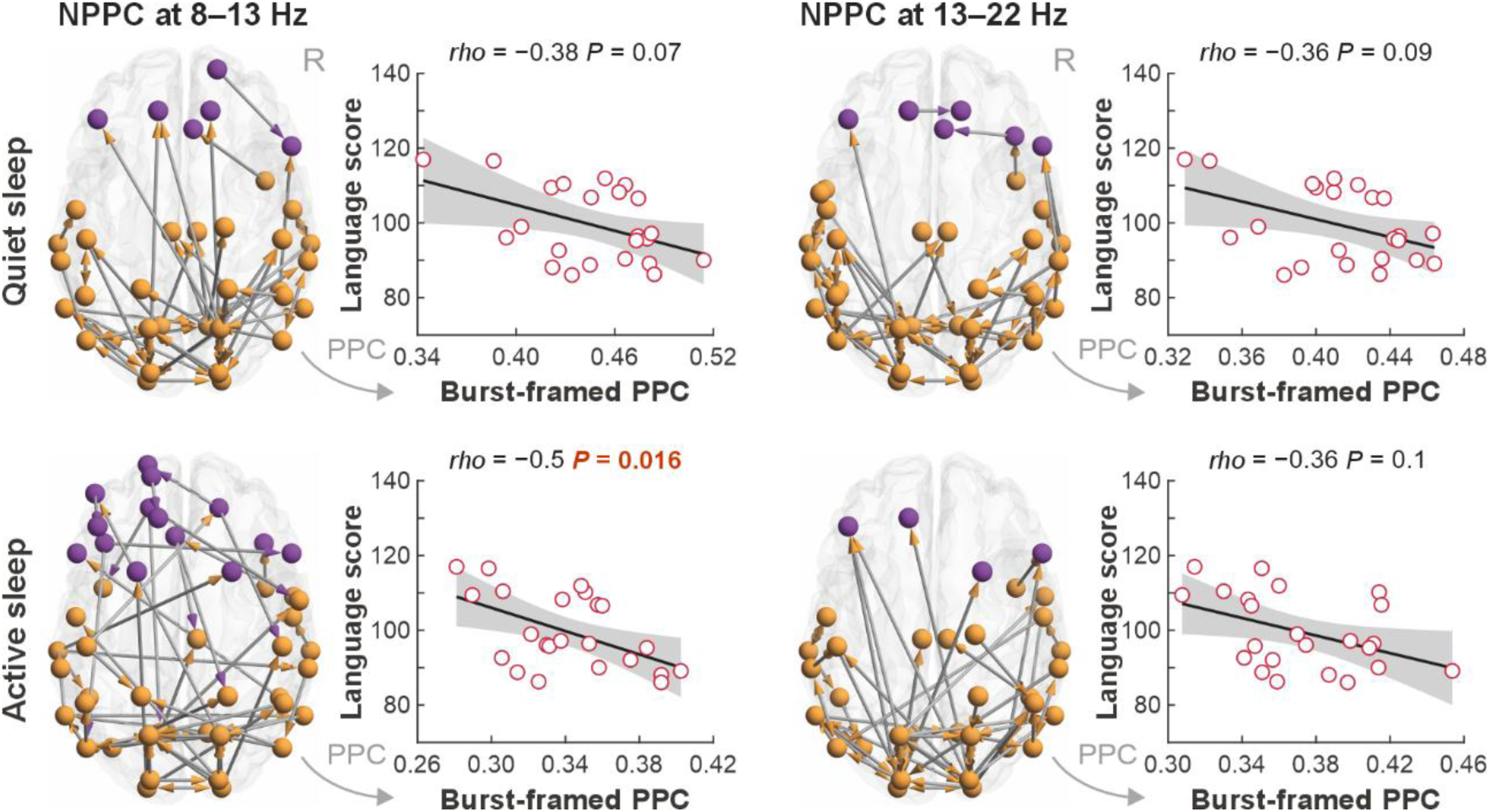
Association between PPC within NPPC networks and neurodevelopment in higher bands. Glass brains show consistent alpha (8–13 Hz) and beta (13–22 Hz) band NPPC networks around term age (37-41 weeks), with quiet sleep on top and active sleep on the bottom. Magenta nodes correspond to the association regions, and orange nodes to the sensorimotor regions. Dots in the scatter plots represent individual infants: mean PPC strength within the network versus corresponding neurodevelopmental scores. Neurodevelopment (cognitive, language, and motor) was assessed using the Bayley Scales at 18 months of age. Correlations were computed using Spearman test. Uncorrected significant correlations are highlighted in red, whereas correlations that passed post hoc correction are shown in bold red. Correlations for cognitive and motor scores for both bands and sleep states were not close to reaching significance (P>0.1).

**Supplementary Figure 8.**
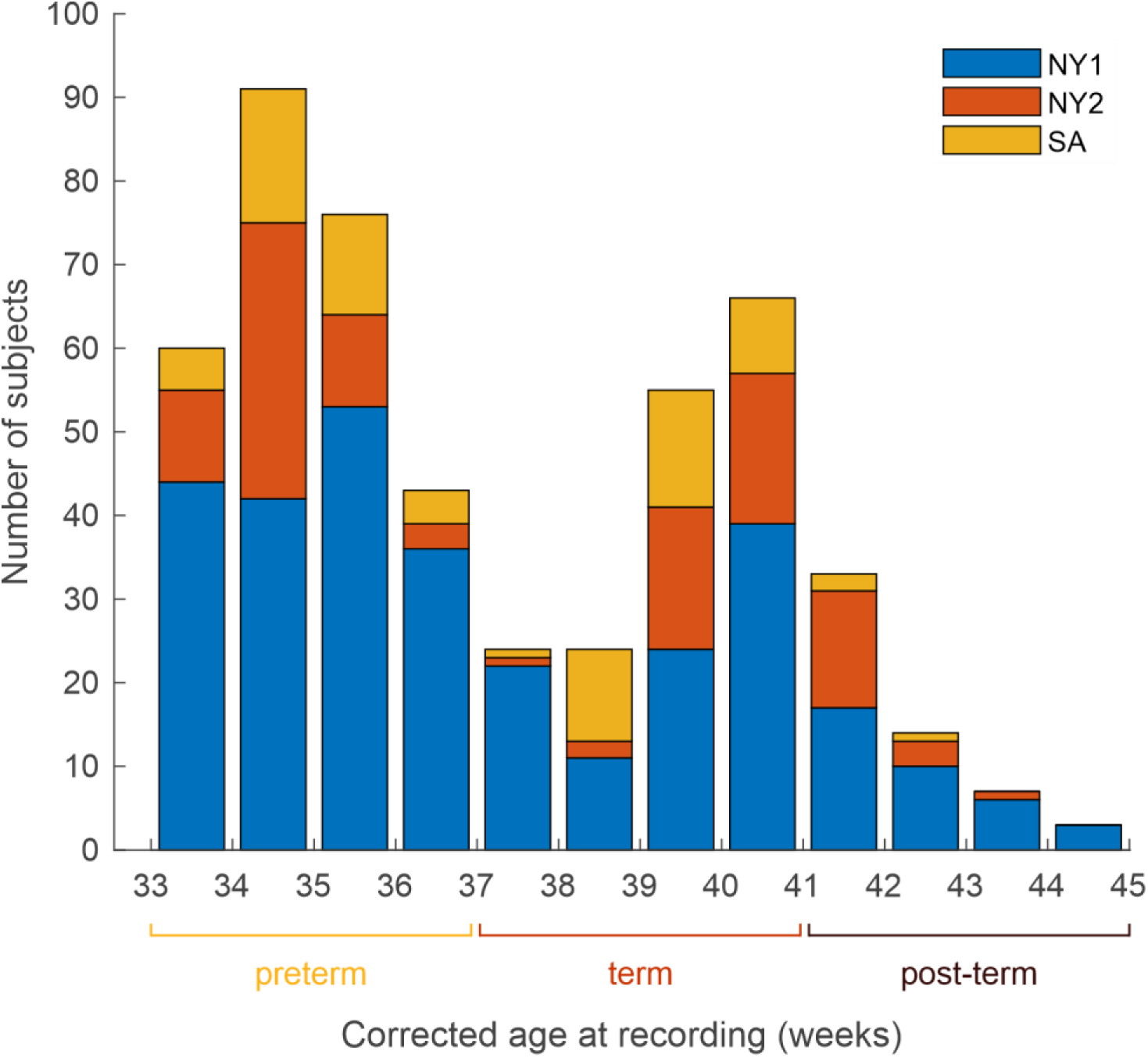
Age distribution of subjects from the three cohorts analyzed in this study.

**Supplementary Figure 9.**
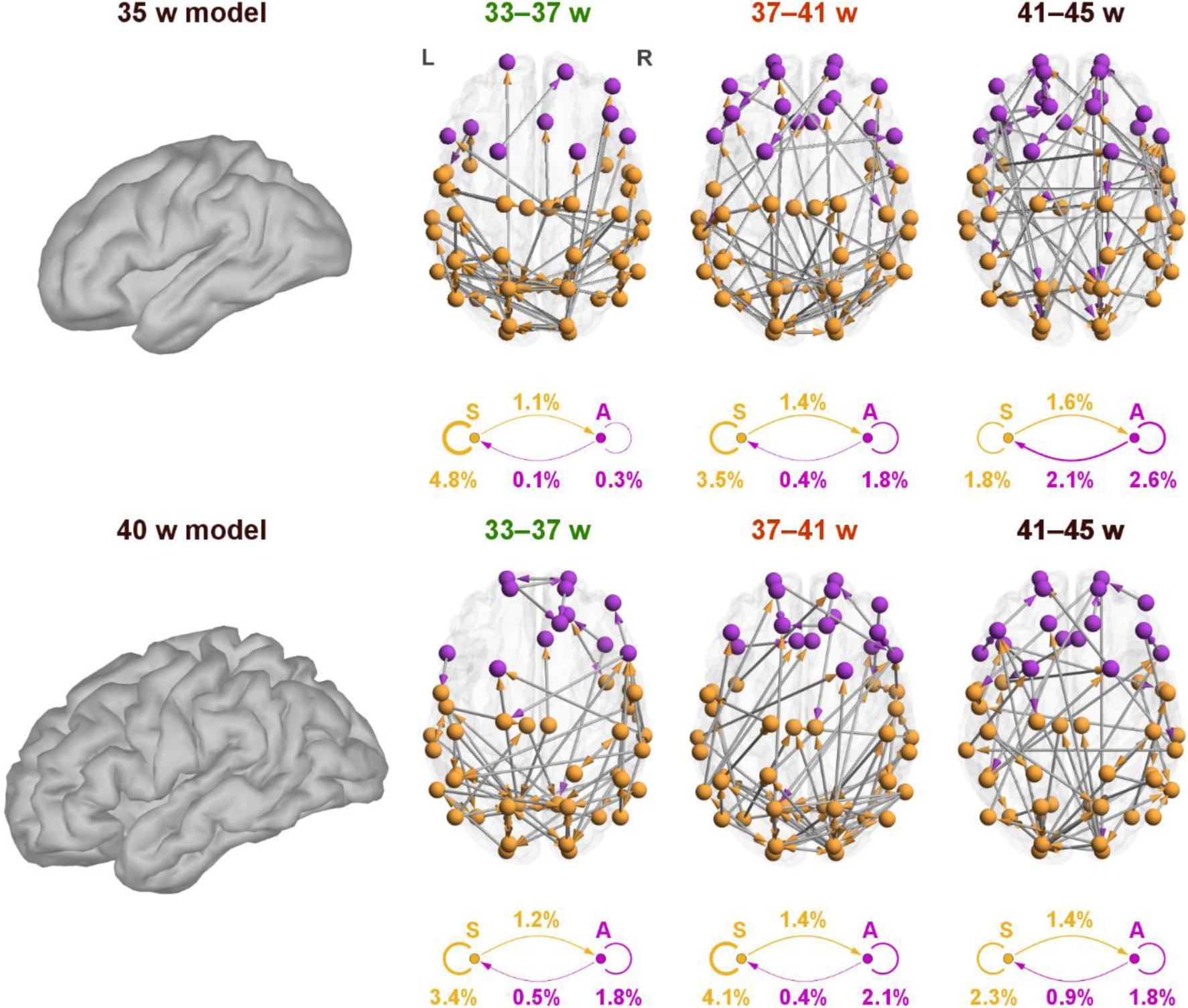
Comparison of head models for smoothed 35-week head model and gyrated 40-week model. Both head models are displayed on the left. NPPC networks for both models in QS theta band are shown on the right.

